# Analysis of costs and strategies for using academic research in a private dental college to develop commercially viable products

**DOI:** 10.1101/2020.06.02.129338

**Authors:** Rampalli Viswa Chandra, Devaraju Rama Raju

## Abstract

**Background & objectives:** The study had two aims. 1) Analysis of research projects done in our institution from 2014-2019 to identify products with a potential for commercialization and 2) To understand the effect of product-development variables on research projects to improve the quality of future commercialization-oriented trials.

**Methods:** 338 clinical trials were grouped into 188 projects under the headings irrigants, diagnostic devices, surgical devices, biomaterials and gels. Trials per project, capital, material costs, labour and the cycle times *per* trial were calculated. To understand the effect these variables, five hypotheses were generated to test whether greater number of trials, successes, higher capital, more investigators per trial and a longer trial duration will result in a product worthy of commercialization.

**Results:** 22 projects had products with a potential for commercialization. Except labour and cycle time (p>0.05), all variables showed significant differences across all projects. Three products were identified as having potential for actual commercialization. It was observed that greater number of trials (χ2=4.6793; p=0.030528) and successes (χ2=20.8134; p<0.00001) in a project along with a higher capital (χ2=12.2662; p=0.000461) will generate a product worthy of commercialization.

**Interpretation & conclusions:** The results seem to suggest that in trials for commercialization, emphasis must be placed on implementing multiple, well-designed clinical trials on a device or product to successfully identify whether it is commercialization-worthy or not. Due attention must be given to the financial aspects of the projects as deficiencies may result in negative impact on the flow and outcomes of a clinical trial.

## INTRODUCTION

Research and development (R&D) have tremendous impact on innovation, and without innovation, prospects for growth in any sector, including Dentistry, may be stunted^1-3^. To conceive and advance an academically sound concept into a marketable product, R&D is heavily dependent on scientific and clinical research^2-4^. High quality academic research and R&D are mutually compatible, and together, will yield a greatest return in the form of new ideas, strategies or products^1-4^.

Frequently functioning under the supervision of a university, dental institutions routinely generate academic research in the form of dissertations and research projects^3-5^. Some of these research projects involve novel therapies, biomaterials or instruments developed by the investigators themselves^2-6^. While results from these studies may or may not be published, after the end-of-study period, most of the ideas or therapies end up in an academic ‘dead-end’ where further development in the form of ‘translational research’^2,5^ is not pursued^3-5^. Bringing a concept from “bench to bedside”^5^ involves hypothesis-driven clinical trials focussing on product development, testing and eventual commercialization^2-4^.

However, product commercialization is a cumbersome process and there are some obvious barriers to the commercialization of an academic idea^2-5^. Academic outcomes^2^ and commercial outcomes^2,5,7^ are quite varied and the paths to derive these outcomes are different as well^2,5-7^. While academic outcomes focus on safety and efficacy of a product^3,5^, commercial outcomes are dependent on how best a product can be modified or evolved for commercial application^4,7^. The ability to identify a commercialization-worthy product requires a thorough understanding on how to identify this ‘gap’^9^ between academic and commercial outcomes^8,9^. Investigators frequently overweigh or under weigh the importance of their own research and the opportunity for the commercialization of their ideas or products^2,5,8,9^. At the same time, all research may not translational in nature and applicable for commercialization^2,5,7^.

One of the first steps in commercialization of a product is the identification and selection of new products and/or ideas^8-10^. In an academic institution which generates a lot of multi-disciplinary data, it can be a daunting task^5,7-11^. More often than not, clinical data is the only record an investigator has access to, and from this, products with clear benefits or outcomes must be identified^7,10^. Every stage of commercialization is also dependent on factors such as the quality of clinical data^8-11^, money spent^11,12^, materials cost ^11,12^ and skilled labour^11^ and the time required to finish a clinical trial or trials^12^ to define a product. Accurate estimation of these variables is essential as these values may be set as a benchmark against which other stages of commercialization are compared to^8-10^. In dentistry at least, there seems to be a paucity of literature regarding identification of commercialization-fit products from various clinical trials and the effect of variables such as capital, material cost, labour and time on commercialization-oriented projects.

In this context, the study had two aims. 1. Analysis of research projects done in our institution within the last five-years to identify products or devices with a potential for commercialization and 2. To understand the effect of product-development variables such as number of trials, capital, material costs, labour and time on clinical trials for defining future strategies to improve quality of research projects in commercialization-oriented projects.

## MATERIALS AND METHODS

All the methodologies and procedures in this study were approved by the Institutional Review Board (SVS-2014-PER-A6).

### Brief profile of the institute

The institute is a post-graduate dental school attached to a medical college and a full-service preclinical research centre capable of toxicology, product assessments, initial product development and exploratory small and large-mammal studies. In 2012, an ‘intellectual property cell’, adhering initially to self-developed guidelines and later amended to conform to the National Intellectual Property Rights Policies, 2016 & 2019^1^, was instituted. Standard operating procedures (SOPs) for conducting clinical trials with a view to promote commercialization in affiliated institutions were developed. Some of the guidelines pertinent to this study were as follows; 1. Within ethical guidelines, novel products, materials or devices developed ‘in-house’ or procured from a source with no commercial interests must be given precedence in clinical trials. 2. Pre-clinical testing such as biocompatibility assays should be performed within the institution whenever possible. 3. For all clinical trials, primary outcomes, which are the variables most relevant to answer the research question must be clearly defined and 4. Data on financial and human resource aspects of clinical trials must be collected from the beginning to the end of a trial in a specified format. All studies from 2013/14 onwards adhered to these guidelines.

### Profile and Growth of R & D projects

In the initial stage, 440 clinical trials (258 dissertations and 182 independent studies) done between 2014-19 in the institution were analyzed. Institutional or self-financed phase II trials on human subjects meeting regulatory standards for ethical research and evaluating novel products, tests or devices with at least two primary outcomes were included in the analysis. Animal studies, *in vitro* investigations and trials on established and commercially available products were excluded.

The primary purpose of every clinical trial was identified and based on its similarity with other trials investigating similar generic products, tests or devices, they were grouped together into ‘projects’ under the following headings: Irrigants, diagnostic devices, surgical devices, biomaterials and gels. 338 clinical trials were grouped into 188 projects under the above headings and the trials required *per* project, money spent (capital/trial) ^11,12^, material cost/trial ^11,12^ (in ₹), skilled labour/trial^11^ and the cycle time/trial^12^ were calculated. Each project yielded a product. Variables in the project were defined as follows^11,13-15,16^; 1. Material cost/trial^⁋^= Production cost+ delivery charges + warranty charges + special equipment charges 2. Cycle time^‡^= time from the beginning to the end of the trial. 3. ^⁑^Labour= a single primary investigator with any number of sub-investigator (or) research assistant. 4. Capital/trial= [Study Costs (Material cost/trail^⁋^ + (administrative staff costs*cycle time^‡^)) + Patient Costs ((procedure cost*subjects) + ((paramedical staff charges + assistant researcher charges)*cycle time^‡^) + (biospecimen processing charges*subjects)) + Labour charges (^⁑^Labour*cycle time^‡^)]. The compounded growth rate (CGR) per year was calculated for 5 years to identify the trend in the number of projects/products fit for commercialization. The CGR (in %)^13^ was calculated as follows; CGR=[(P_final_/P_begin_)^1/t^-1] *100 where P_begin_ and P_final_ are the number of products at the beginning and end of the year of the year and ‘t’ is time in years.

### Hypothesis generation

To understand the effect of variables such as trials *per* project, capital, material cost, labour and cycle time on defining future strategies to improve quality of clinical trials in commercialization-oriented projects, the following hypotheses were generated; H1: Greater number of trials in a project will improve its product-generation prospects. H2: Greater successes (k) in clinical trials in a project will generate a product worthy of commercialization. H3: A higher capital and H4: man-force is needed to generate a product that can be commercialized. H5: A higher cycle-time is required to generate a commercialization worthy-product.

### Setting a cutoff in to identify products with potential for commercialization

Probability of k-success events in Bernoulli trials^14,17-22^ was calculated for all trials and individual trails to determine whether a product is suitable for commercialization or not. Briefly it was done as follows; all trials were considered as Bernoulli experiments under the assumption that 1. Each trial was independent. 2. Each trial results in one of two possible outcomes, success (S) or failure (F) and 3. The probability of S remains constant (p=0.5) from trial-to-trial and is denoted by p and 1-p is the constant probability. For all trials (n), the number of successful (k) and non-successful (n-k) outcomes were calculated. A trial was considered as successful if all of the primary outcomes were fulfilled. The probability of success for all trials (binomial distribution) was calculated from the formula P_o_ (k successes in n trials) = (^n^ _k_) ^pk^(1-p) ^n-k^. For individual trials, n=number of outcomes, k=number of successful outcomes and n-k were unsuccessful outcomes with p=k/n. The results for all the trials (P_o_) and individual trials (P_i_) were obtained. As the sample size was large, normal approximation to the binomial was done for P_o_ and were approximated to the normal model with parameters µ = np and σ = √np (1 − p). The top 10% values were calculated through Z_10%_= (X_1_-µ)/σ to obtain X_1_ which was the cutoff score for products with the potential for commercialization. For individual studies with P_i_ ≥x_1,_ the overall trial scores were calculated separately again and the top 10% values were obtained to obtain a cutoff score (X_2_) to identify products for actual commercialization [Z_10%_= (X_2_-µ)/σ]. Project and device-wise probability score averages at initial analysis and identification of products with potential for commercialization and actual commercialization are described as P_k,_ P_1_ and P_2_ respectively.

### Identification of products with the potential for commercialization

Projects with X_1_≥0.15^17-22^ were assumed as having products with potential for commercialization^21,22^. Descriptive data such as P_1,_ average capital, material cost and labour to develop a product along with time taken for each trial (cycle time) were calculated for these projects again.

### Identification of products for actual commercialization

Projects with X_2_≥0.20 ^17-22^, were assumed as having products with potential for actual commercialization. P_2_, number of trials, subjects, total costs, material costs, total labour and cycle time for the entire project were calculated as mean weighted averages.

### Determinants of Commercialization

H1 vs >1 trial/project, H2 vs >2 successes/project, H3 vs > 3,00,000 ₹/trial, H4 vs >4 individuals /trial and H5 vs >12 months/trial were assessed to examine the relation between the hypotheses and the variables associated with them.

### Statistical Analysis

Data was analyzed by using Prism8^®^ (GraphPad Software, La Jolla, USA). Intragroup comparison was performed by using ANOVA followed by multiple comparisons using Bonferroni correction. Unpaired t-test was used for intergroup comparisons. A chi-square test of independence was performed to examine the relation between the hypotheses and the variables associated with them. A p≤0.001 was considered as highly significant, p≤0.05 as significant and p>0.05 as non-significant. A panel of five investigators worked separately from the main study investigators to analyse the data and perform the required calculations.

## RESULTS

### Profile and Growth of R & D projects

338 institutional or self-financed trials from 2014 to 2019 on human subjects evaluating novel products, tests or devices (phase II) were grouped into 188 projects. *Table I* summarizes the distribution of variables such as P_k_-values, number of projects, trials per project, capital, material cost, labour and cycle time under the headings: Irrigants, diagnostic devices, surgical devices, biomaterials and gels. A significant to highly significant distribution *(p=0.001)* was seen for number of projects, P_k_ scores *(p=0.02)*, trials per project *(p=0.04)*, capital and material costs across the headings. Surgical devices had least number of projects (4 out of 188); however, this category also had the highest numbers for all the other variables. The opposite was observed for projects under irrigants. Highest P_k_ score was observed for projects in diagnostic devices group. CGR per year for projects was not constant; rather there were yearly variations. From 2014-19 (79 projects) to 2018-19 (188 projects), the CGR growth over five years was 19.23% per year.

**Table I:** Products identified from 188 projects (338 clinical trials; 2014-19) and the distribution of variables under them.

| Products | No. of Projects | P<sub>k</sub> /Project | Trials per project | Capital/trial (₹) | Material Cost (₹)/trial | Average Labour /trial | Cycle time/trial (Months) |
| --- | --- | --- | --- | --- | --- | --- | --- |
| Irrigants | 92 | 0.13±0.07 | 1.08±0.19 | 2,19,405±1,20,005 | 26,038±20,831 | 3±2 | 9±1.2 |
| Diagnostic devices | 12 | 0.17±0.12 | 1.66±0.70 | 11,84,806±1,72,083 | 4,96,474±2,27,179 | 7±3 | 14±3.8 |
| Surgical devices | 4 | 0.16±0.09 | 6.25±1.25 | 22,68,778±9,25,021 | 5,80,327±2,76,087 | 11±6 | 16±5.2 |
| Biomaterials | 36 | 0.15±0.08 | 3.33±1.65 | 7,01,464±2,84,389 | 29,359±22,491 | 3±2 | 11±4.0 |
| Gels | 44 | 0.14±0.04 | 1.70±0.82 | 3,64,770±1,57,818 | 73,965±34,416 | 3±1 | 9±4.1 |
| *F-Value* | 62.81 | 6.32 | 66.92 | 143.78 | 98.21 | 1.56 | 17.23 |
| *p value* | 0.001** | 0.02* | 0.04* | 0.001** | 0.001** | 0.34‡ | 0.20‡ |

P<sub>k</sub> =k successes in n trials ‡Non-significant \*Significant \*\*Highly significant

### Identification of projects with the potential for commercialization

X_1_≥0.15 was the chosen cut-off value to identify products with potential for commercialization. The initial pool of 188 research projects narrowed down to 22 commercialization-focussed projects with 59 trials in total. *Table II* summarizes the distribution of variables such as P_1_-values, number of projects, trials per project, capital, material cost, labour and cycle time under project headings. A trend similar to previous observations made before the cut-off was seen. Previously insignificant, a significant distribution *(p=0.0126)* was seen for labour/trial across all projects. Higher number of projects were seen for biomaterial and irrigant groups.

**Table II:** Identification of projects identified as having a potential for commercialization after applying the cut-off (X_1_≥0.15; 22 projects, 59 trials).

| Products | Number | P<sub>1</sub><br>/Project | Trials/<br>Project | Capital/ trial<br>(₹) | Material Cost//<br>trial (₹) | Labour/trial<br>(individuals) | Cycle<br>time/<br>trial<br>(Months) |
| --- | --- | --- | --- | --- | --- | --- | --- |
| Irrigants | 8 | 0.17±0.04 | 1.21±0.01 | 3,53,147±28,680 | 40,756±9,358 | 4±1 | 9±0.8 |
| Diagnostic<br>devices | 1 | 0.24±0.06 | 2.01±0.24 | 13,75,984±3,86,960 | 7,41,089±1,58,195 | 8±2 | 16±3.2 |
| Surgical<br>devices | 1 | 0.19±0.07 | 7.29±0.22 | 26,40,643±4,82,285 | 7,93,393±1,43,176 | 13±4 | 18±4.3 |
| Biomaterials | 9 | 0.20±0.04 | 4.22±0.69 | 9,12,794±97,136 | 93,794±18,902 | 4±2 | 12±2.7 |
| Gels | 3 | 0.15±0.01 | 2.02±0.41 | 4,92,097±39,247 | 1,16,910±17,510 | 3±1 | 10±3.4 |
| *F-Value*<br>*p value* | 6.98<br>0.01* | 1.22<br>0.012* | 23.81<br>0.02* | 66.92<br>0.04* | 67.82<br>0.00011** | 6.43<br>0.0126* | 1.53<br>0.2282‡ |

P<sub>1</sub> =k successes in n trials ‡Non-significant \*Significant \*\*Highly significant

*Table III* summarizes the comparison of all variables before and after the identification of commercialization-worthy projects through the application of cut-off of probability of success scores. The measures for labour and cycle time remained unaffected *(p>0.05)*. Trials/project for irrigants *(p=0.1)* and surgical devices *(p=0.09)* did not show significant differences as well. P_k_ vs P_1_ values for gels remain unaffected. The remaining variables showed significant to highly significant differences across all projects.

**Table III:** Comparison of all variables before and after the identification of commercialization-worthy projects through the application of X_1_ cut-off of probability of success scores.

| Type of the products | Number of projects | $P_k$ vs $P_1$ /Project | Trials/ Project | Capital/ trial (₹) | Material Cost/ trial (₹) | Labour/trial (individuals) | Cycle time/ trial (Months) |
| --- | --- | --- | --- | --- | --- | --- | --- |
| *Irrigants* | $\leq 0.001^{**}$ | $\leq 0.001^{**}$ | 0.1‡ | $\leq 0.001^{**}$ | $\leq 0.001^{**}$ | 0.09‡ | 0.7‡ |
| *Diagnostic devices* | $\leq 0.001^{**}$ | $\leq 0.001^{**}$ | 0.003* | 0.03* | 0.05* | 0.07‡ | 0.09‡ |
| *Surgical devices* | $\leq 0.001^{**}$ | $\leq 0.001^{**}$ | 0.09‡ | 0.004* | $\leq 0.001^{**}$ | 0.06‡ | 0.07‡ |
| *Biomaterials* | $\leq 0.001^{**}$ | $\leq 0.001^{**}$ | 0.04* | $\leq 0.001^{**}$ | $\leq 0.001^{**}$ | 0.3‡ | 0.1‡ |
| *Gels* | $\leq 0.001^{**}$ | 0.2‡ | 0.05* | $\leq 0.001^{**}$ | $\leq 0.001^{**}$ | 0.1‡ | 0.09‡ |

$P_k$ & $P_1$ =k successes in n trials ‡Non-significant \*Significant \*\*Highly significant

### Identification of products for actual commercialization

Projects with X_2_≥0.20^11,20^, were assumed as having products with potential for actual commercialization. Three products were identified as having a potential for commercialization. They are 1. A low-cost oral cancer detection device *(Product 1)* 2. A butyrate inactivated recombinant human bone morphogenetic protein-2 *(rhBMP-2)* gel for bone regeneration *(Product 2)*. 3. basic fibroblast growth factor (bFGF) impregnated collagen membranes for soft-tissue regeneration *(Product 3) (Figure I). Table IV* summarizes the products identified for commercialization and the variables under them. Total project cost was higher for product-3 whereas material costs were higher for product-2. Both the products also had more trials per project (4/project). Cycle time, labour required and subjects under the project were maximum for product-1.

**Figure I:**
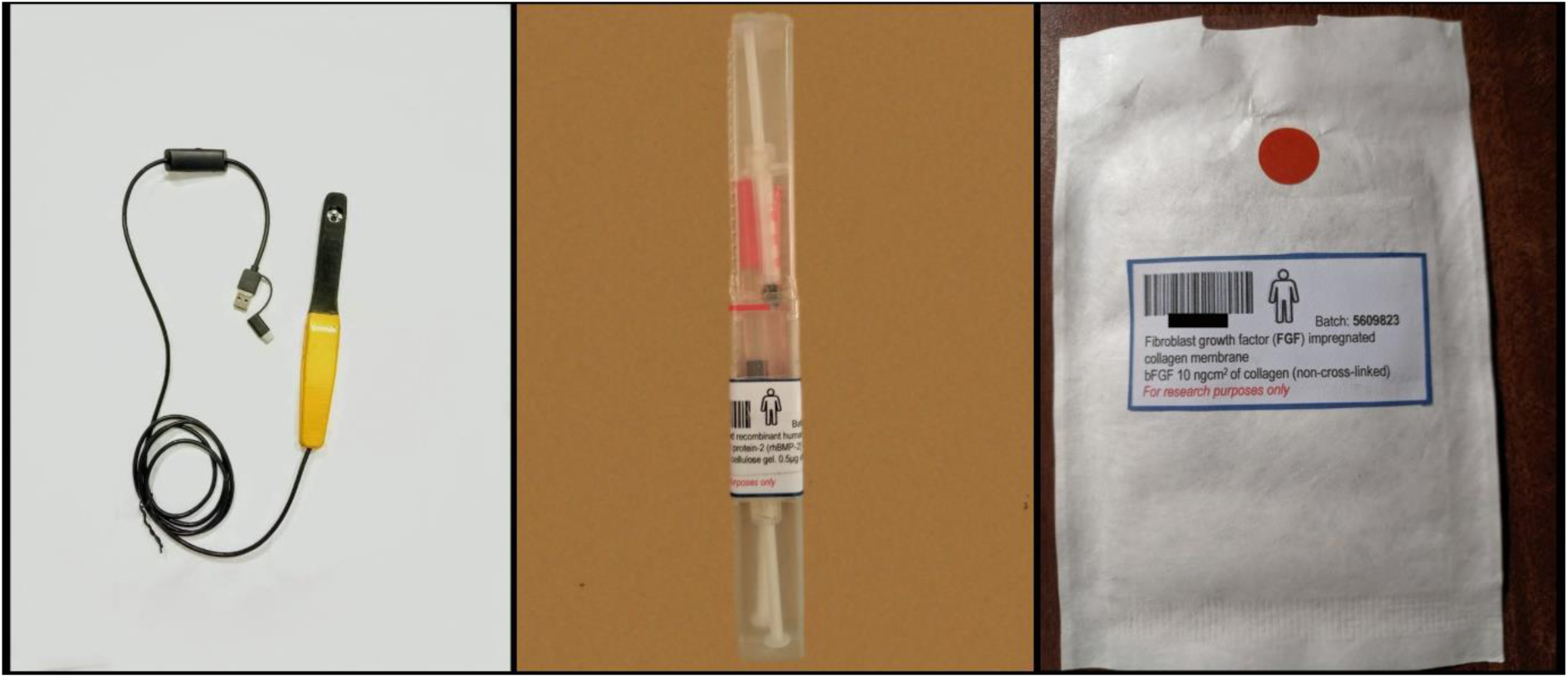
Three products were identified as having a potential for commercialization. They are 1. A low-cost oral cancer detection device *(left)* 2. A butyrate inactivated recombinant human bone morphogenetic protein-2 (rhBMP-2) gel for bone regeneration *(middle)* and 3. basic fibroblast growth factor (bFGF) impregnated collagen membranes for soft-tissue regeneration *(right).*

**Table IV:** Products identified for final commercialization and the distribution of variables under them (X_2_≥0.20).

| **Products** | **P<sub>2</sub>** | **Trials/**<br>**Project** | **Subjects** | **Total**<br>**Cost**<br>**(₹)** | **Material**<br>**Cost (₹)** | **Total**<br>**Labour**<br>**(individuals)** | **Cycle**<br>**time**<br>**(Months)** |
| --- | --- | --- | --- | --- | --- | --- | --- |
| **Oral cancer**<br>**detection**<br>**device** | 0.28 | 2 | 187<br>(3) | 40,54,743<br>(1,93,439) | 14,47,459<br>(1,92,774) | 15<br>(1) | 18<br>(2) |
| **RhBMP-2 gel** | 0.23 | 4 | 118<br>(2) | 42,63,356<br>(1,57,063) | 19,06,649<br>(1,19,401) | 14<br>(2) | 13<br>(2) |
| **bFGF**<br>**impregnated**<br>**collagen**<br>**membranes** | 0.23 | 4 | 97<br>(3) | 44,50,534<br>(1,49,968) | 15,93,025<br>(1,73,441) | 15<br>(1) | 14<br>(2) |

<sup>a)</sup> Reported numbers represent weighted averages and SD (in parenthesis) of five observations rounded off to the nearest integer. P<sub>2</sub> =k successes in n trials.

### Determinants of Commercialization

On comparing hypotheses with their associated variables, it appears that greater number of trials (χ2=4.6793; p=0.030528) and successes (χ2=20.8134; p<0.00001) in a project along with a higher capital (χ2=12.2662; p=0.000461) will generate a product worthy of commercialization. The number of investigators/trial and the trial duration seem to have no effect on the outcomes of a commercialization *(Table V)*.

**Table V:**
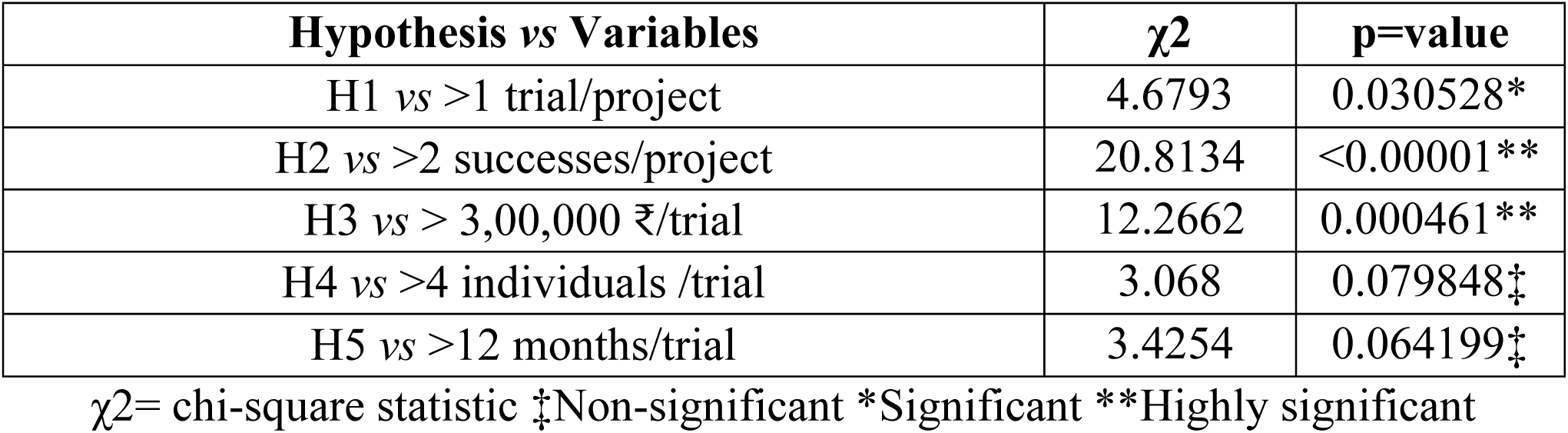
Comparison of hypotheses with their associated variables; H1: Greater number of trials H2: Greater successes (k) in clinical trials H3: Higher capital H4: Higher man-force and H5: A higher cycle-time

## DISCUSSION

The private sector essentially controls and dominates product development and commercialization^2-4,6^; Academic institutes were traditionally thought of as centres that contribute to innovative patents, but do not pursue development or commercialization of the ideas or devices behind the patents^3,6^. Institutions including dental schools generate a lot of research, but there is a need to ensure that the output is constant and in tune with the principles of product development and commercialization^2-4^. A steady increase in projects and trials is an essential part of product development and commercialization^9,10^ and a 10% CGR over the five-year period is an indicator of healthy growth in commercialization-worthy projects^9,10,13^. We observed a CGR of 19% per year over a period of five years.

In this study, greater number of trials and successes of those trials (k) were significantly associated with products worthy of commercialization. Except for gels, highly significant difference was seen for trials/project and P_k_ vs P_1_ values before and after the identification of commercialization-worthy projects. Higher number of commercialization based-projects were seen for biomaterial (9) and irrigant groups (8), but irrigant-based projects had fewer number of trials (1.21±0.01 vs 4.22±0.69 trials/project) than biomaterial projects. Though there was only a single project under diagnostic devices, its success in trials was the highest (P_2_=0.28). These findings can be related to the products selected for final commercialisation; one diagnostic device and two biomaterials. Institutes may have to start or evaluate a huge number of clinical trials under multiple projects in order to achieve one commercial success and the prediction of success becomes a difficult task. Investigators have sought to group or increase the number of trials per project more efficiently through various means to assess products during pre-commercialization trials^23,24^. Studies have successfully utilized web-based tools^23^ or have created repositories^24^ for similar trials under a project to focus primarily on introducing new products or ideas. At the same time, a higher rate of success of clinical trials in a project can be achieved by careful product identification^9^, adequate personnel training^10,13,19,25^ and production of high-quality data^25^.

Product development and commercialization are money-driven and as in any form or research, capital plays an important role in sustaining clinical trials^8,10,11^; we observed that projects with a higher capital and material costs tend to generate a product worthy of commercialization. Significant to highly significant differences were seen for capital before and after the X_1_ cut-off. Irrigants had lower capital and material costs whereas diagnostic and surgical devices needed higher capital and material costs. The capital required was still lower when compared to grants received/requested to develop or test similar devices. For example, the low-cost oral cancer detection device and rhBMP-2 gel that we seek to commercialize utilized a maximum capital of 40,54,743₹ and 42,63,356₹ respectively; studies evaluating a similar products have quoted a capital 3 to 18 times higher including development costs^26,27^. Over contract research organizations (CRO), academic institutes are uniquely placed to offer an advantage of reduced development and commercialization costs because of the following reasons; 1. Availability of “third-stream” academicians^28^ proficient in R&D activities can lower man-power costs by eliminating certain positions such as trial manager etc^3,25,28^. 2. Presence of a ready organisational support that can deal with various aspects of research such as subject recruitment, allotment, interventions, data collection and analysis, all of which, can translate into substantial savings when commercialization is being pursued^1-4^.

Material costs, also called ‘external costs’, is often a significant portion of capital required to run a trials-to-commercialization study^10,11,15,28,29^. It includes procurement, purification, manufacturing and packaging costs of a product along with any additional expenses incurred towards achieving regulatory approval and patenting^15,29^. In R&D, around 30% of the budget is spent on product development whereas the remainder is on clinical trials^8,9,11,12,15^. The material costs were 35 to 44% of the total capital for the three products identified for commercialization. However, it is difficult to compare these values with benchmarks from earlier studies as all material costs are factored into the final pricing of the material and end up in a clinical trial as “per-patient” or “per-site” costs^15,27,29,30^; one study^15^ reported that the ‘clinical procedure costs’ consumed about 22.32% of the overall costs. Most of the projects in this study procured materials either in bulk or small batches which made it easy for us to calculate material costs *via* invoicing.

The number of investigators (labour) per trial and the trial duration seem to have no effect on the outcomes of a commercialization. Development of products and technologies is highly dependent on the availability of skilled manpower^8,10,13^. In a study on 207 collaborative projects, there were no significant differences in terms of personnel and duration as well^3^. It was also observed that trail duration is generally recorded accurately as the number of patients recruited, therapy delivered and maintenance is dependent on this variable^12,15,23,25,30^. We recognized only three roles; a single primary investigator (PI) with any number of sub-investigators or research assistants. Roles such as ‘clinical trials manager’ or ‘clinical trials coordinator’ were curtailed as their frequent intervention is known to generate ‘overpowered’^15^ and unusable data^30^. Though the number of subjects and number of trials for the final three products were different, the study duration and the number of personnel were almost similar. As most of the studies were academic dissertations and short projects done by post-graduate residents, adequate manpower in form of post-graduates and their supervisors were always available and the study duration would be closer to academic standards adjusted up or down depending on the trial^6,8,10,23,25^.

This study has some limitations worth noting. Prediction of products which can be successfully commercialized is based on complex frameworks and prediction models which are often of the single-use type and is limited to one product^5,8,9,18,19,21^. We chose a relatively simple tool unconventionally in the form of probability of k-success events in Bernoulli trials^14,17-22^. This metric is versatile and can be used for sample size estimation^14^, clinical trial designs^17^, stochastic predictions^18^, financial analysis^19^, diagnosis^20^, biomarker assessment^21^ and in decision making^22^. Though we have focussed on phase II trials, a commercialization-worthy product is identified at the preclinical stage itself and the transition of this product in various phases of clinical trials (phases I-IV) is rigorously evaluated in a time frame that can range from years to a decade^5,9,10^. We were limited by the study-period of our primary investigators, most of whom were post-graduates. The nature of clinical trial was given little significance as assessment of the product was done at a project level rather than a clinical-trial level. We feel this is not a disconcerting limitation as evaluation of too many trial dependent factors may result in an unpredictable behaviour of the metric and variables analyzed^14,18,20^.

## CONCLUSION

To conclude, we had analyzed research projects done in our institution within the last five-years to identify products or devices with the potential for commercialization and sought to understand the effect of product-development variables to define future strategies to improve quality of clinical trials in commercialization-oriented projects. Three products were identified and commercialization of these products is being actively pursued and one of the devices has already been patented. The results seem to suggest that in trials for commercialization, emphasis must be placed on implementing multiple well-designed clinical trials on a device or product to successfully identify whether it is commercialization-worthy or not. Due attention must be given to the financial aspects of the projects as deficiencies may result in negative impact on the flow and outcomes of a clinical trial.

## Supporting information

author form- NON ACADEMIC DOCUMENT

author form- NON ACADEMIC DOCUMENT

## Notes

Source of financial support: This study is supported by a grant (DST/NSTMIS/05/222/2017-18) under the National Science & Technology Management Information System (NSTMIS), Department of Science and Technology, Government of India.

### Competing Interest Statement

This study is supported by a grant (DST/NSTMIS/05/222/2017-18) under the National Science & Technology Management Information System (NSTMIS), Department of Science and Technology, Government of India. Dr Rama Raju holds a patent on a low-cost oral cancer detection device. Dr R Viswa Chandra has patents pending on a butyrate inactivated recombinant human bone morphogenetic protein-2 (rhBMP-2) gel and basic fibroblast growth factor (bFGF) impregnated collagen membranes.

### Summary of Updates

We have removed Sr and II from the authors names respectively. Apologize for the inconvenience but we are not comfortable using those.

